## Supplementary_Information for "Voxelated bioprinting of modular double-network bio-ink droplets"


**The PDF file includes:**

Supplementary Figs. 1 to 9

**Other Supplementary Materials for this manuscript include the following:**

Supplementary Movies 1 to 6

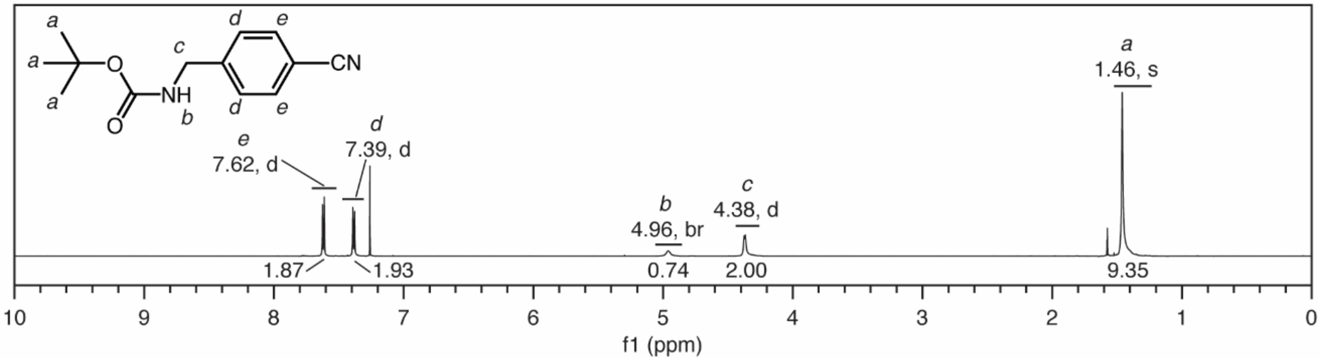


**Supplementary Fig. 1. $^{1}$H NMR spectrum of *tert*-butyl *N*-(4-cyanobenzyl)carbamate**.

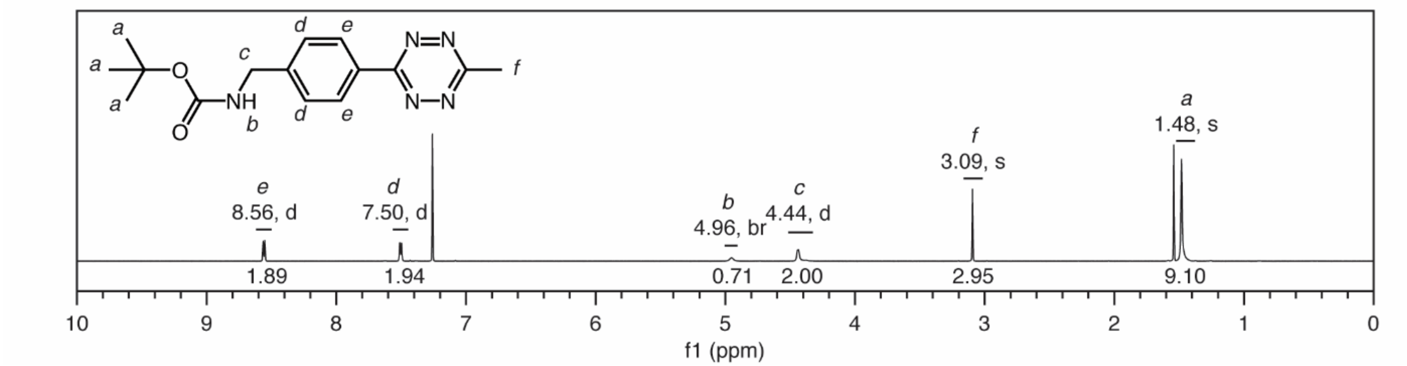


**Supplementary Fig. 2.** $^{1}$**H NMR of** ***tert*****-butyl** ***N*****-[4-(6-methyl-1,2,4,5-tetrazin-3-yl)benzyl]carbamate**.

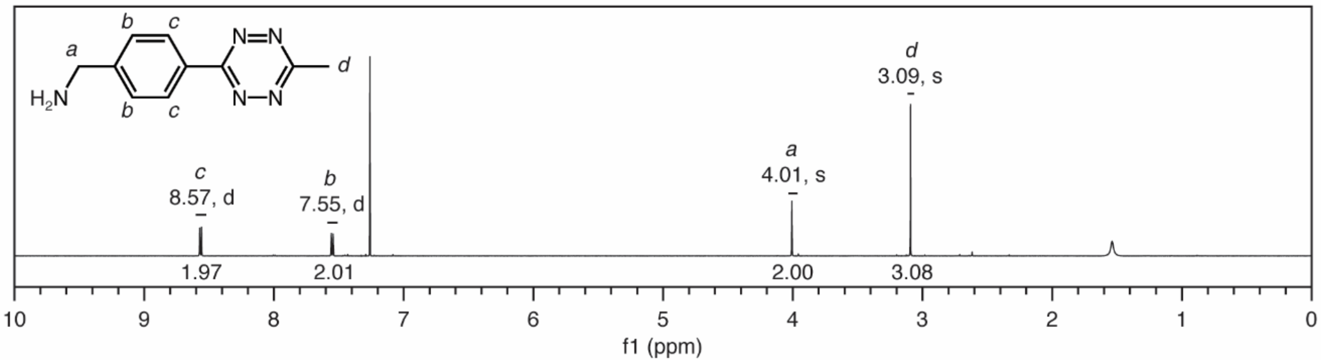


**Supplementary Fig. 3. $^1$H NMR of [4-(6-methyl-1,2,4,5-tetrazin-3-yl) phenyl]methanamine**.

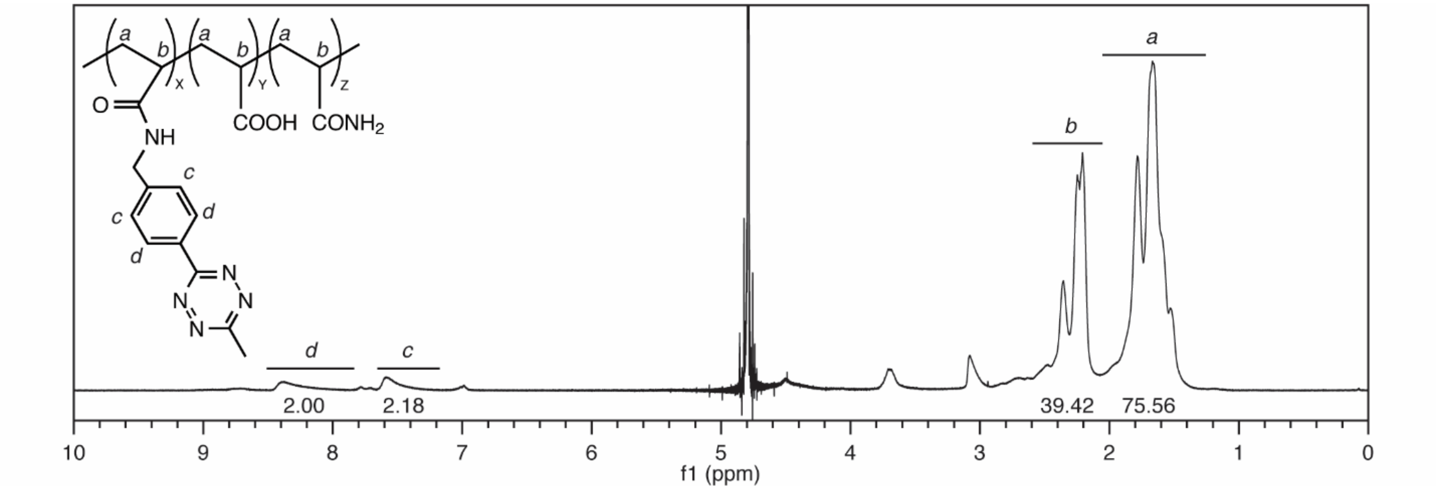


**Supplementary Fig. 4. $^{1}$H NMR of tetrazine modified poly(acrylamide-*co*-acrylic acid)**.

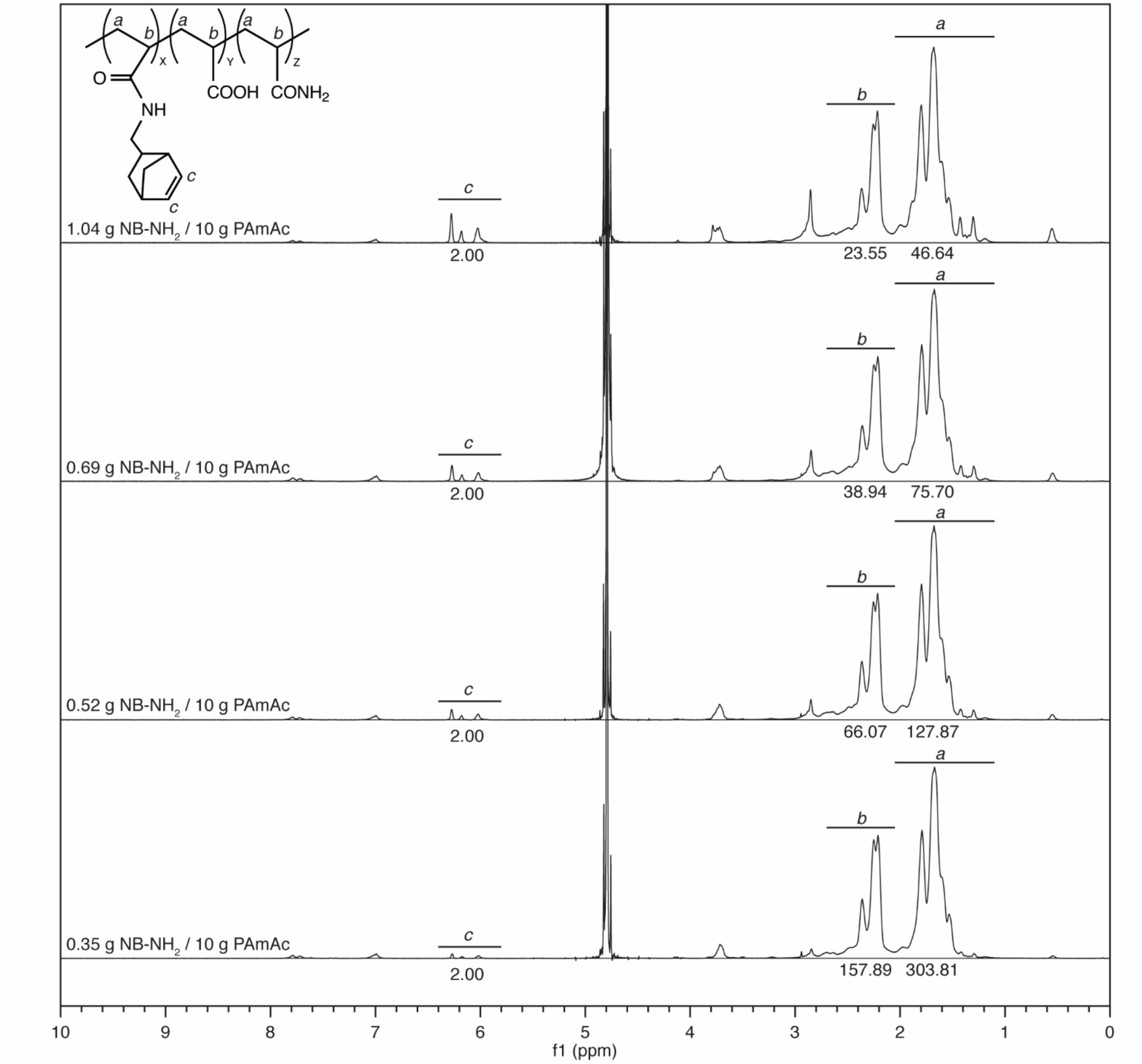


**Supplementary Fig. 5. $^1$H NMR of norbornene modified poly(acrylamide-*co*-acrylic acid)**.

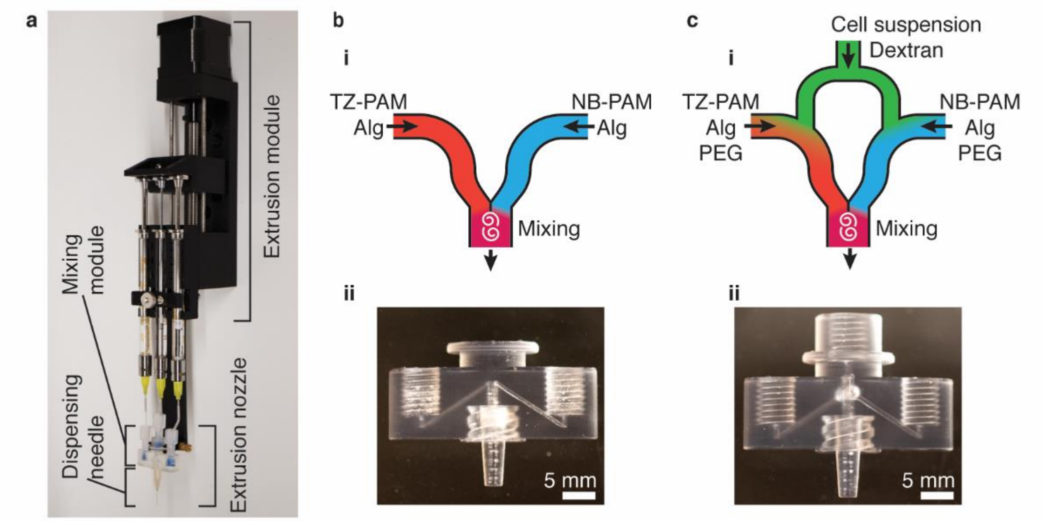


**Supplementary Fig. 6. Setup of the DASP 2.0**. (**a**) Assembly of the extrusion module, syringes, and extrusion nozzle. (**b**) (i) Schematic and (ii) photograph of the dual-inlet mixing module. (**c**) (i) Schematic and (ii) photograph of the triple-inlet mixing module.

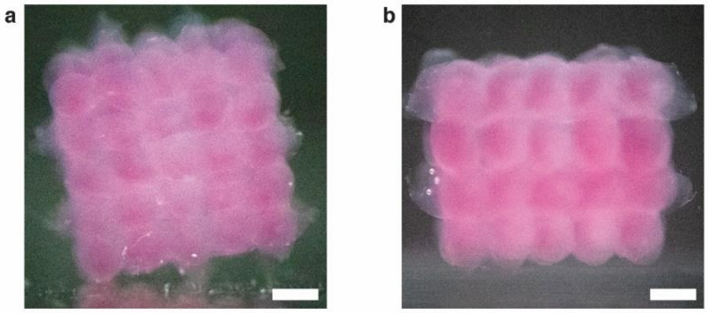


**Supplementary Fig. 7. A DASP printed 3D lattice.** The 5×5×4 lattice consists of 100 interconnected yet distinguishable hydrogel particles. (**a**) and (**b**) display the top and lateral view, respectively. Scale bar, 1 mm.

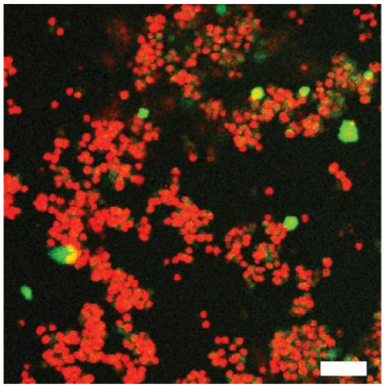

**Supplementary Fig. 8. A representative fluorescence confocal microscopy image from live/dead assay of MIN6 cells**. The cells with a density of $2\times10^7$/mL are incubated in DMEM with 10% w/v polyacrylamide for 3h and subsequently suspended to cell culture media to reach a final density of $1.2\times10^5$/mL for cell recovery. The image is taken 12 h after the recovery. Scale bar, 50 μm.

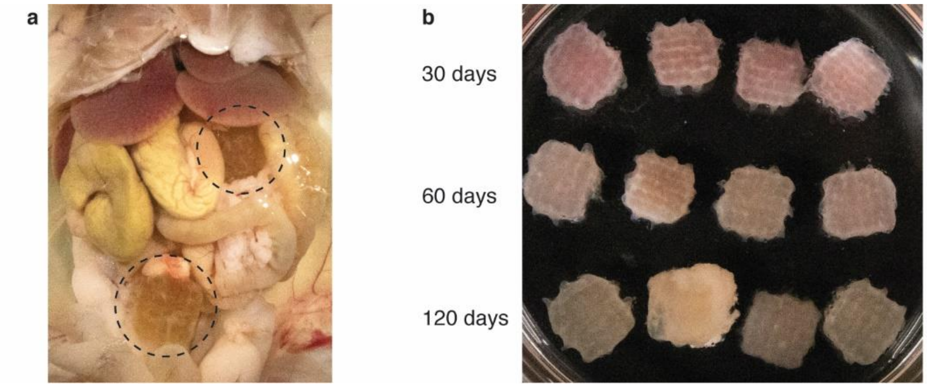


**Supplementary Fig. 9. Representative photograph of the retrieved scaffolds**. (**a**) A photograph of the scaffolds made of DN hydrogel located near the liver in the abdominal cavity (dashed circles). (**b**) A photograph of the all scaffolds retrieved at 30, 60 and 120 days after transplantation.

**Supplementary Movie 1.** Tensile tests of dog-bone shape samples of double-network hydrogels.

**Supplementary Movie 2.** Mixing a pair of viscoelastic bio-inks using a dual-inlet print nozzle.

**Supplementary Movie 3.** Manipulation of a hollow sphere printed by DASP 2.0.

**Supplementary Movie 4.** A DASP 2.0 printed hollow sphere cut into two pieces.

**Supplementary Movie 5.** Manipulation of a DASP 2.0 printed gyroid made of double-network hydrogel.

**Supplementary Movie 6.** Cyclic compression test of a gyroid made of double-network hydrogel.
